# Treatment of infection-induced vascular pathologies is protective against persistent rough morphotype *Mycobacterium abscessus* infection in zebrafish

**DOI:** 10.1101/2021.11.23.469750

**Authors:** Julia Y Kam, Kathryn Wright, Warwick J Britton, Stefan H Oehlers

## Abstract

*Mycobacterium abscessus* infections are of increasing global prevalence and are often difficult to treat due to complex antibiotic resistance profiles. While there are similarities between the pathogenesis of *M. abscessus* and tuberculous mycobacteria, including granuloma formation and stromal remodeling, there are distinct molecular differences at the host-pathogen interface. Here we have used a zebrafish-*M. abscessus* model and host-directed therapies that were previously identified in the zebrafish-*M. marinum* model to identify potential host-directed therapies against *M. abscessus* infection. We find efficacy of anti-angiogenic and vascular normalizing therapies against rough *M. abscessus* infection, but no effect of anti-platelet drugs.

## Background

*Mycobacterium abscessus* is a species of non-tuberculous mycobacteria (NTM) that is found in abundance within a range of environments [1, 2]. There has been a rise in the global incidence of *M. abscessus* infection in recent years, with the bacteria frequently identified in severe respiratory, skin and mucosal infections [3]. Pulmonary *M. abscessus* infection is especially prevalent in patients with pre-existing chronic lung diseases, such as cystic fibrosis and bronchiectasis [4, 5]. Disease management and treatment is difficult as *M. abscessus* is extremely resistant to antibiotic and anti-tuberculous drug treatments [6-8]. Preclinical *M. abscessus* virulence is further complicated by the existence of rough (R) and smooth (S) colony morphotypes which is dependent on the absence or presence of surface-exposed glycopeptidolipids (GPLs), respectively, which are reduced or lost in R variants by transcriptional downregulation or loss of function mutations [9-14]. As the morphotypes display divergent modes of pathogenesis, it is likely that they evoke different host responses to infection.

Host-directed therapies (HDTs) are a treatment approach aimed at modulating the host immune response to a pathogen rather than targeting the pathogen directly. Unlike antibiotics, the infectious agent is not able to mutate resistance of the HDT target [15]. For example, pazopanib, metformin, and imatinib have been shown to control mycobacterial infection through inhibition of angiogenic signaling, improved bactericidal responses, and modulation of apoptosis respectively [16-18].

There are presently few HDTs identified with efficacy against *M. abscessus*. A recent study showed that AR-12, a celecoxib derivative, has direct and host-mediated antimicrobial effects in a neutropenic mouse model of *M. abscessus* infection through unidentified mechanisms [19]. Another study has found that sirtuin-3 (SIRT3), a mitochondrial protein deacetylase, is an important host factor in murine and zebrafish control of *M. abscessus* [20]. Infected mice and zebrafish treated with the SIRT3 agonist resveratrol displayed significantly reduced bacterial load and histopathological damage [20].

*M. abscessus* infection is cleared by immunocompetent mice within weeks, with immunodeficient knockout strains or corticosteroid treatment required to sustain infection in mice [21-23]. Thus, it can be challenging to delineate the effects of HDT treatment from the natural clearance of infection in an immunologically intact mouse. We recently demonstrated that adult zebrafish maintain a high bacterial burden along with granuloma formation and recruitment of cellular immunity for weeks after initial *M. abscessus* infection [24]. This suggests a zebrafish model may be better suited for the study of persistent *M. abscessus* infection and testing the efficacy of potential HDTs.

We have identified HDTs inhibiting aspects of the infection-induced vascular pathologies caused by *M. marinum* in zebrafish. These include aspirin and other anti-platelet drugs that inhibit infection-induced thrombocytosis, pazopanib that blocks infection-induced angiogenesis, and the experimental drug, AKB-9785 that reverses infection-induced vascular permeability [18, 25, 26]. Whether targeting these host pathways during *M. abscessus* infection has therapeutic benefits remains unknown. Therefore, we investigated whether these drugs may be potential HDTs to treat *M. abscessus* infection.

## Methods

### Zebrafish lines and handling

Zebrafish breeding stock were held in the Centenary Institute under Sydney Local Health District Animal Welfare Committee (SLHD AWC) approval 17-036. Experiments on adult zebrafish were carried out in accordance with SLHD AWC approvals 17-037 and 20-029.

Zebrafish embryos were produced by natural spawning and raised in E3 media at 28°C. Wild type zebrafish were the AB strain background and *Tg(kdrl:gfp)*^*s843*^ fish were used to visualize vasculature.

### Handling of *M. abscessus* strains and infection

Rough (R) and smooth (S) variants of *M. abscessus* strain CIP104536^T^ were grown at 37°C in Middlebrook 7H9 broth supplemented with 10% Oleic acid/Albumin/Dextrose/Catalase (OADC) enrichment and 0.05% Tween 80 or on Middlebrook 7H10 agar containing 10% OADC (7H10 OADC). Recombinant *M. abscessus* strains expressing tdTomato or Wasabi were grown in the presence of 300 µg/ml hygromycin [9, 27]. Homogenous bacterial suspensions for injection were prepared as previously reported [28], embryos were infected by microinjection into the neural tube with approximately 1,000 CFU and adult fish were infected with 20,000 to 50,000 CFU by intraperitoneal injection.

### Live Imaging

Embryos were anesthetized in tricaine (MS222 160 µg/ml, Sigma) and mounted in 3% methylcellulose for live imaging using a Leica M205FA fluorescent microscope.

### Drug Treatment of Infected Adult Zebrafish

Infected zebrafish were treated at 1 week post infection (wpi) (rough *M. abscessus*) or 3 wpi (smooth *M. abscessus*) with final concentrations of 50 μM AKB-9785, 10 μM aspirin, 1 μM pazopanib, or 10 μM tirofiban. Drugs were refreshed every second day for 1 week (rough *M. abscessus*) or 2 weeks (smooth *M. abscessus*).

### Bacterial Recovery

Animals were euthanised by tricaine anaesthetic overdose (>300□μg/ml) and rinsed in PBS. Whole carcasses were mechanically homogenised and serially diluted in PBS. Homogenates were plated onto 7H10 OADC supplemented with 300 μg/ml hygromycin. Plates were grown for at least 3 days at 37°C.

### Histology

Animals were subjected to cryosectioning as previously described [29]. Briefly, euthanasia was performed by tricaine anaesthetic overdose and specimens were fixed for 2-4 days in 10% neutral buffered formalin at 4°C. Specimens were then rinsed in PBS, incubated overnight in 30% sucrose, followed by an overnight incubation in a 50/50 solution of 30% sucrose and Tissue-Tek O.C.T. compound (OCT). A final overnight incubation in OCT was performed prior to freezing at -80°C. Cryosectioning was performed to produce 20 µm thick sections. Sections were post-fixed in 10% neutral buffered formalin and rinsed in PBS prior to further processing. Slides for fluorescent imaging were mounted with coverslips using Fluoromount G containing DAPI.

Slides were imaged on a Leica DM6000B microscope. Consecutive images for granuloma census were taken 200 μm apart through the entire abdominal cavity.

### Image analysis

All image processing and analysis were performed using Fiji-ImageJ using fluorescent pixel counts or measurement as previously described [29].

### Statistics

All statistical analysis and graphing were performed using GraphPad Prism 9 (GraphPad Software). Statistical tests were carried out as indicated, error bars represent standard deviation, and P-values are supplied in figures.

## Results

### Rough *M. abscessus* is susceptible to vascular-targeted HDTs

We first examined whether infection of zebrafish with *M. abscessus* induces a host angiogenic response as seen in *M. marinum* infection [18]. The vascular reporter line *Tg(kdrl:gfp)*^*s843*^, where blood endothelial cells are labelled with GFP, was utilized to visualize infection-associated angiogenesis. Embryos were infected with rough or smooth *M. abscessus* (*M. abscessus* R or *M. abscessus* S) and imaged at 6 dpi to assess ectopic blood vessel growth. While we observed infection-induced angiogenesis in *M. abscessus* R-infected embryos, we did not observe an angiogenic response to *M. abscessus* S (Figure 1A).

**Figure 1:**
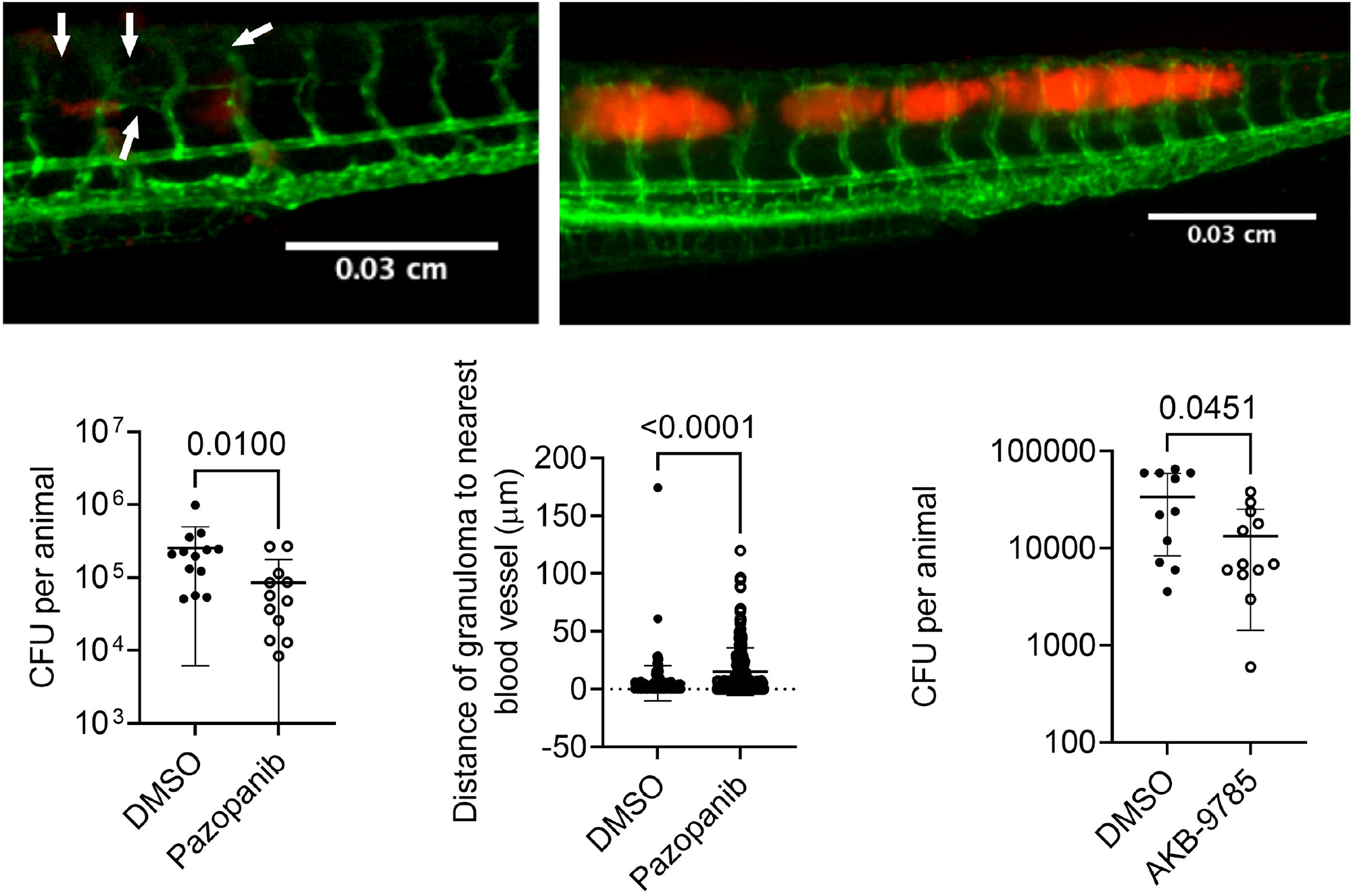
Vascular targeted therapies reduce rough *M. abscessus* burden. (A) Representative images of rough and smooth *M. abscessus* infection-induced angiogenesis (white arrows) in 6 dpi *Tg(kdrl:gfp)*^*s843*^ embryos. (B) Quantification of rough *M. abscessus* burden in adult zebrafish treated with pazopanib. Each data point represents a single fish. Data is representative of three experiments. (C) Quantification of minimum distance between *M. abscessus* granuloma and nearest blood vessel. Each data point represents a single granuloma, total animals analyzed DMSO: 4, Pazopanib: 4. (D) Quantification of rough *M. abscessus* burden in adult zebrafish treated with AKB-9785. Each data point represents a single fish. Data are representative of two experiments. Statistical testing by Student’s t test D, Mann Whitney U test B, C.

To further assess whether inhibiting angiogenesis would affect *M. abscessus* growth during chronic infection we treated infected adult zebrafish with the VEGFR inhibitor pazopanib [18]. Treatment with pazopanib reduced *M. abscessus* R, but not *M. abscessus* S, burden (Figure 1B and Supplementary Figure 1A). To validate the effectiveness of pazopanib treatment against infection-induced angiogenesis we quantified the distance between granulomas and their closest blood vessel (Supplementary Figure 1B). Treatment with pazopanib increased the distance between *M. abscessus* R granulomas and their closest blood vessel (Figure 1C).

To investigate the role of vascular permeability on *M. abscessus* infection we treated infected adult zebrafish with the VE-PTP inhibitor AKB-9785 to induce vascular normalization [25]. Treatment with AKB-9785 reduced the burden of *M. abscessus* R (Figure 1D), but not *M. abscessus* S (Supplementary Figure 1C).

### *M. abscessus* infection is not sensitive to platelet-targeted therapies

To assess whether inhibition of infection-induced thrombocyte activation is therapeutic against *M. abscessus* infection we treated adult zebrafish with aspirin or tirofiban [26]. Aspirin treatment did not impact the bacterial burden of *M. abscessus* R or *M. abscessus* S (Figure 2A and 2B). Similarly, we did not observe any effect of tirofiban treatment on either morphotype of *M. abscessus* (Figure 2C and 2D).

**Figure 2:**
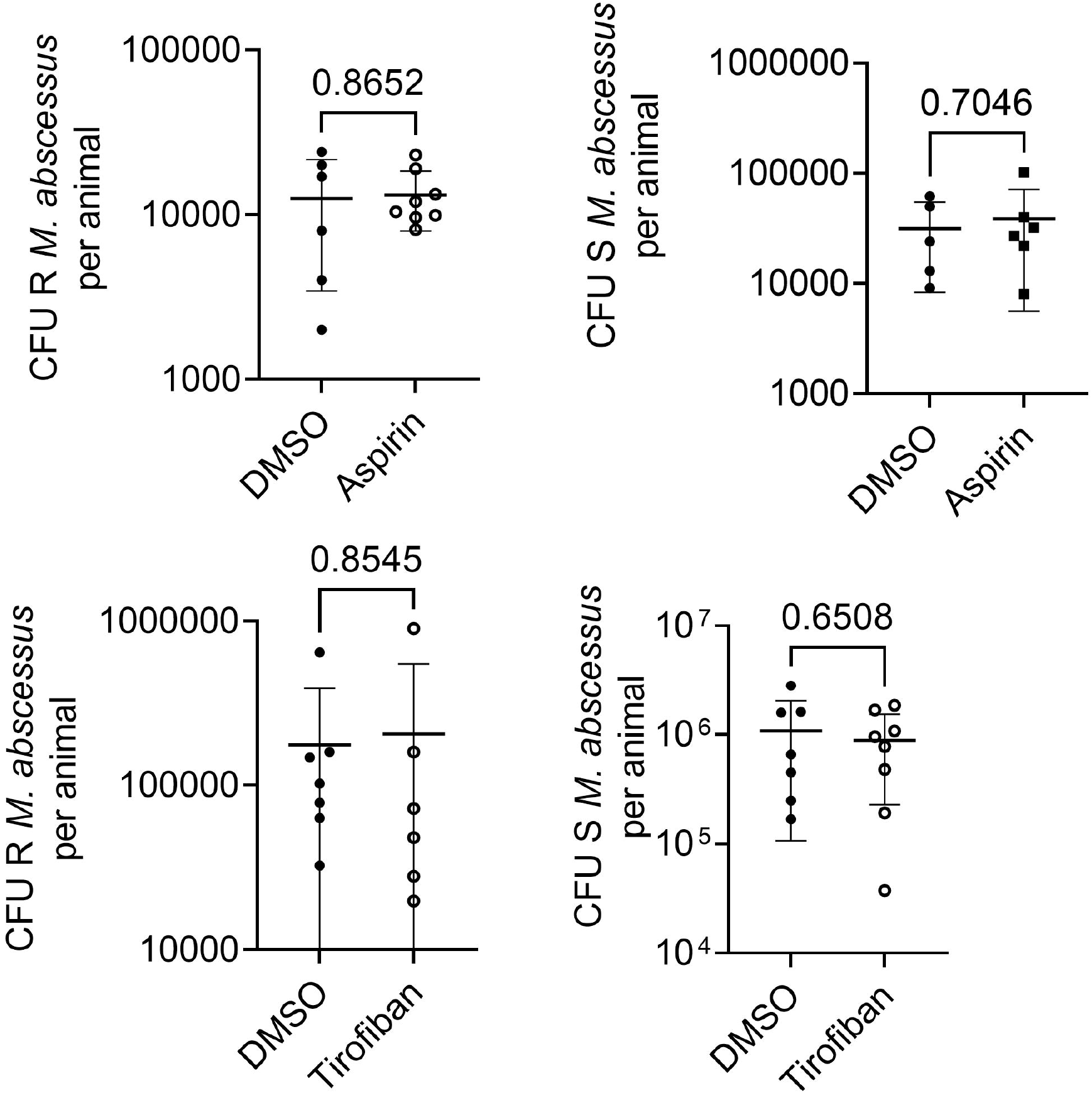
Anti-platelet drugs do not affect *M. abscessus* burden. (A) Quantification of rough *M. abscessus* burden in adult zebrafish treated with aspirin. Each data point represents a single fish. Data are representative of 3 experiments. (B) Quantification of smooth *M. abscessus* burden in adult zebrafish treated with aspirin. Each data point represents a single fish. Data are representative of 3 experiments. (C) Quantification of rough *M. abscessus* burden in adult zebrafish treated with tirofiban. Each data point represents a single fish. (D) Quantification of smooth *M. abscessus* burden in adult zebrafish treated with tirofiban. Each data point represents a single fish. Statistical testing by Student’s t test.

## Discussion

We sought to explore potential HDTs with the zebrafish-*M. abscessus* model, using identified HDTs that we had previously validated in the zebrafish-*M. marinum* model of tuberculosis. We show that like tuberculous mycobacterial infections, inhibition of infection-induced angiogenesis and vascular permeability restrict the growth of rough *M. abscessus*. Conversely, vascular targeted therapies do not affect the growth of smooth *M. abscessus*. Furthermore, inhibition of thrombocyte activation with aspirin or tirofiban did not affect control of either *M. abscessus* morphotype.

*M. tuberculosis* and *M. marinum* have been shown to drive vascularization and vascular permeability around their granulomas [18, 30]. Here, we found that, like tuberculous mycobacteria, rough *M. abscessus* induces host angiogenesis and that this angiogenesis can be targeted to reduce rough *M. abscessus* burden. However, we did not observe infection-induced angiogenesis in smooth *M. abscessus*-infected zebrafish embryos and there was no effect of anti-angiogenic therapy on smooth *M. abscessus* burden, suggesting that morphotype-specific differences in granuloma pathogenesis may drive this difference. We have observed that rough *M. abscessus* forms larger necrotic granulomas more rapidly than smooth *M. abscessus* in adult zebrafish infected with equivalent doses [31]. We hypothesize the granulomatous nature of rough *M. abscessus* growth exposes the granuloma-resident mycobacteria to more hypoxic conditions than smooth *M. abscessus*. It will be interesting to determine if the loss of cell surface GPLs during smooth to rough transition increases production of VEGF by infected macrophages by revealing similar cell wall components to other mycobacteria [32, 33].

Elevated platelet activation has been observed in cystic fibrosis patients, which may be aggravated by respiratory infections [34, 35]. Our finding that anti-platelet drugs did not affect *M. abscessus* burden is different to our findings in *M. marinum* infection, where treatment with aspirin or tirofiban reduce *M. marinum* burden [26]. We speculate that *M. abscessus* may not activate zebrafish hemostasis or cause thrombocytosis. While this could be further investigated by infecting hemostasis-reporting zebrafish with *M. abscessus* to visualize infection-induced hemostasis, further investigation in standard zebrafish is unlikely to aid in the identification of anti-hemostatic HDT strategies. Rather we hypothesize that a stronger phenotype may be observed in a model with elevated baseline platelet activation caused by depletion of Cystic fibrosis transmembrane conductance regulator.

A major implication of the observations made during this study was that the host immune system behaves differently against rough *M. abscessus*, compared to both smooth *M. abscessus* and *M. marinum*. Our current understanding of the interaction between tuberculous mycobacteria and the host immune system may have limited application to rough *M. abscessus* pathology, and therefore future studies should focus on investigating the unique host-*M. abscessus* dynamics *in vivo* to reveal potential therapeutic targets.

## Footnote page

### Meetings where the information has previously been presented

None.

## Acknowledgements

The authors thank Drs Laurent Kremer and Matt Johansen for supplying *M. abscessus* strains and assistance with handling, members of the Tuberculosis Research Program at the Centenary Institute for discussion of the manuscript, Dr Angela Fountaine of the BioImaging Facility and Sydney Cytometry at Centenary Institute for technical assistance with imaging, and Aerpio Therapeutics for AKB-9785.

## Figure Legends

**Supplementary Figure 1**

(A) Quantification of smooth *M. abscessus* burden in adult zebrafish treated with pazopanib. Each data point represents CFU recovered from a single fish. Data are representative of two experiments. (B) Example image of a necrotic granuloma containing rough *M. abscessus*-TdTomato (orange) in a DAPI-stained (blue) section from *Tg(kdrl:gfp)*^*s843*^ adult zebrafish (green blood vessels). Arrow indicates closest blood vessel to edge of granuloma. (C) Quantification of smooth *M. abscessus* burden in adult zebrafish treated with AKB-9785. Each data point represents CFU recovered from a single fish. Statistical testing by Student’s t test.

## References

1. Thomson R, Tolson C, Sidjabat H, Huygens F, Hargreaves M. Mycobacterium abscessus isolated from municipal water - a potential source of human infection. BMC infectious diseases 2013; 13:241.

2. Burgess W, Margolis A, Gibbs S, Duarte RS, Jackson M. Disinfectant Susceptibility Profiling of Glutaraldehyde-Resistant Nontuberculous Mycobacteria. Infection control and hospital epidemiology 2017; 38:784–91.

3. Johansen MD, Herrmann J-L, Kremer L. Non-tuberculous mycobacteria and the rise of Mycobacterium abscessus. Nat Rev Microbiol 2020; 18:392–407.

4. Esther CR, Esserman DA, Gilligan P, Kerr A, Noone PG. Chronic Mycobacterium abscessus infection and lung function decline in cystic fibrosis. Journal of Cystic Fibrosis 2010; 9:117–23.

5. Chu H, Zhao L, Xiao H, et al. Prevalence of nontuberculous mycobacteria in patients with bronchiectasis: a meta-analysis. Arch Med Sci 2014; 10:661–8.

6. Nessar R, Cambau E, Reyrat JM, Murray A, Gicquel B. Mycobacterium abscessus: a new antibiotic nightmare. Journal of Antimicrobial Chemotherapy 2012; 67:810–8.

7. Jeon K, Kwon OJ, Lee NY, et al. Antibiotic Treatment of Mycobacterium abscessus Lung Disease. American Journal of Respiratory and Critical Care Medicine 2009; 180:896–902.

8. Chopra S, Matsuyama K, Hutson C, Madrid P. Identification of antimicrobial activity among FDA-approved drugs for combating Mycobacterium abscessus and Mycobacterium chelonae. Journal of Antimicrobial Chemotherapy 2011; 66:1533–6.

9. Bernut A, Herrmann JL, Kissa K, et al. Mycobacterium abscessus cording prevents phagocytosis and promotes abscess formation. Proc Natl Acad Sci U S A 2014; 111:E943–52.

10. Howard ST, Rhoades E, Recht J, et al. Spontaneous reversion of Mycobacterium abscessus from a smooth to a rough morphotype is associated with reduced expression of glycopeptidolipid and reacquisition of an invasive phenotype. Microbiology 2006; 152:1581–90.

11. Medjahed H, Reyrat JM. Construction of Mycobacterium abscessus defined glycopeptidolipid mutants: comparison of genetic tools. Appl Environ Microbiol 2009; 75:1331–8.

12. Gutierrez AV, Viljoen A, Ghigo E, Herrmann JL, Kremer L. Glycopeptidolipids, a Double-Edged Sword of the Mycobacterium abscessus Complex. Front Microbiol 2018; 9:1145.

13. Pawlik A, Garnier G, Orgeur M, et al. Identification and characterization of the genetic changes responsible for the characteristic smooth-to-rough morphotype alterations of clinically persistent Mycobacterium abscessus. Mol Microbiol 2013; 90:612–29.

14. Le Moigne V, Bernut A, Cortes M, et al. Lsr2 Is an Important Determinant of Intracellular Growth and Virulence in Mycobacterium abscessus. Front Microbiol 2019; 10:905.

15. Kilinç G, Saris A, Ottenhoff THM, Haks MC. Host-directed therapy to combat mycobacterial infections. Immunological Reviews.

16. Napier RJ, Norris BA, Swimm A, et al. Low Doses of Imatinib Induce Myelopoiesis and Enhance Host Anti-microbial Immunity. PLOS Pathogens 2015; 11:e1004770.

17. Singhal A, Jie L, Kumar P, et al. Metformin as adjunct antituberculosis therapy. Science translational medicine 2014; 6:263ra159.

18. Oehlers SH, Cronan MR, Scott NR, et al. Interception of host angiogenic signalling limits mycobacterial growth. Nature 2015; 517:612–5.

19. Zhang S, Zou Y, Guo Q, et al. AR-12 Exhibits Direct and Host-Targeted Antibacterial Activity toward Mycobacterium abscessus. Antimicrobial Agents and Chemotherapy 2020; 64:e00236–20.

20. Kim YJ, Lee S-H, Jeon SM, et al. Sirtuin 3 is essential for host defense against Mycobacterium abscessus infection through regulation of mitochondrial homeostasis. Virulence 2020; 11:1225–39.

21. Bernut A, Le Moigne V, Lesne T, Lutfalla G, Herrmann J-L, Kremer L. In Vivo Assessment of Drug Efficacy against Mycobacterium abscessus Using the Embryonic Zebrafish Test System. Antimicrobial Agents and Chemotherapy 2014; 58:4054.

22. Lerat I, Cambau E, Roth dit Bettoni R, et al. In Vivo Evaluation of Antibiotic Activity Against Mycobacterium abscessus. The Journal of Infectious Diseases 2013; 209:905–12.

23. Maggioncalda EC, Story-Roller E, Mylius J, Illei P, Basaraba RJ, Lamichhane G. A mouse model of pulmonary Mycobacteroides abscessus infection. Scientific reports 2020; 10:3690-.

24. Hortle E, Kam JY, Krogman E, et al. Rough and smooth variant Mycobacterium abscessus infections are differentially controlled by host immunity during chronic infection of adult zebrafish. bioRxiv 2020:856948.

25. Oehlers SH, Cronan MR, Beerman RW, et al. Infection-induced vascular permeability aids mycobacterial growth. J Infect Dis 2017; 215:813–7.

26. Hortle E, Johnson KE, Johansen MD, et al. Thrombocyte Inhibition Restores Protective Immunity to Mycobacterial Infection in Zebrafish. J Infect Dis 2019; 220:524–34.

27. Bernut A, Nguyen-Chi M, Halloum I, Herrmann JL, Lutfalla G, Kremer L. Mycobacterium abscessus-Induced Granuloma Formation Is Strictly Dependent on TNF Signaling and Neutrophil Trafficking. PLoS Pathog 2016; 12:e1005986.

28. Bernut A, Dupont C, Sahuquet A, Herrmann JL, Lutfalla G, Kremer L. Deciphering and Imaging Pathogenesis and Cording of Mycobacterium abscessus in Zebrafish Embryos. J Vis Exp 2015.

29. Cheng T, Kam JY, Johansen MD, Oehlers SH. High content analysis of granuloma histology and neutrophilic inflammation in adult zebrafish infected with Mycobacterium marinum. Micron 2020; 129:102782.

30. Datta M, Via LE, Kamoun WS, et al. Anti-vascular endothelial growth factor treatment normalizes tuberculosis granuloma vasculature and improves small molecule delivery. Proc Natl Acad Sci U S A 2015; 112:1827–32.

31. Hortle E, Kam J, Krogman E, et al. Rough and smooth variant Mycobacterium abscessus infections are differentially controlled by host immunity during chronic infection of adult zebrafish. bioRxiv 2021.

32. Rhoades ER, Archambault AS, Greendyke R, Hsu FF, Streeter C, Byrd TF. Mycobacterium abscessus Glycopeptidolipids mask underlying cell wall phosphatidyl-myo-inositol mannosides blocking induction of human macrophage TNF-alpha by preventing interaction with TLR2. J Immunol 2009; 183:1997–2007.

33. Walton EM, Cronan MR, Cambier CJ, et al. Cyclopropane Modification of Trehalose Dimycolate Drives Granuloma Angiogenesis and Mycobacterial Growth through Vegf Signaling. Cell Host Microbe 2018; 24:514–25 e6.

34. O’Sullivan BP, Linden MD, Frelinger AL, 3rd, et al. Platelet activation in cystic fibrosis. Blood 2005; 105:4635–41.

35. Ortiz-Munoz G, Yu MA, Lefrancais E, et al. Cystic fibrosis transmembrane conductance regulator dysfunction in platelets drives lung hyperinflammation. J Clin Invest 2020; 130:2041–53.

