## Supplementary figures and images for "Treatment of infection-induced vascular pathologies is protective against persistent rough morphotype *Mycobacterium abscessus* infection in zebrafish"

### Supplementary Figure 1

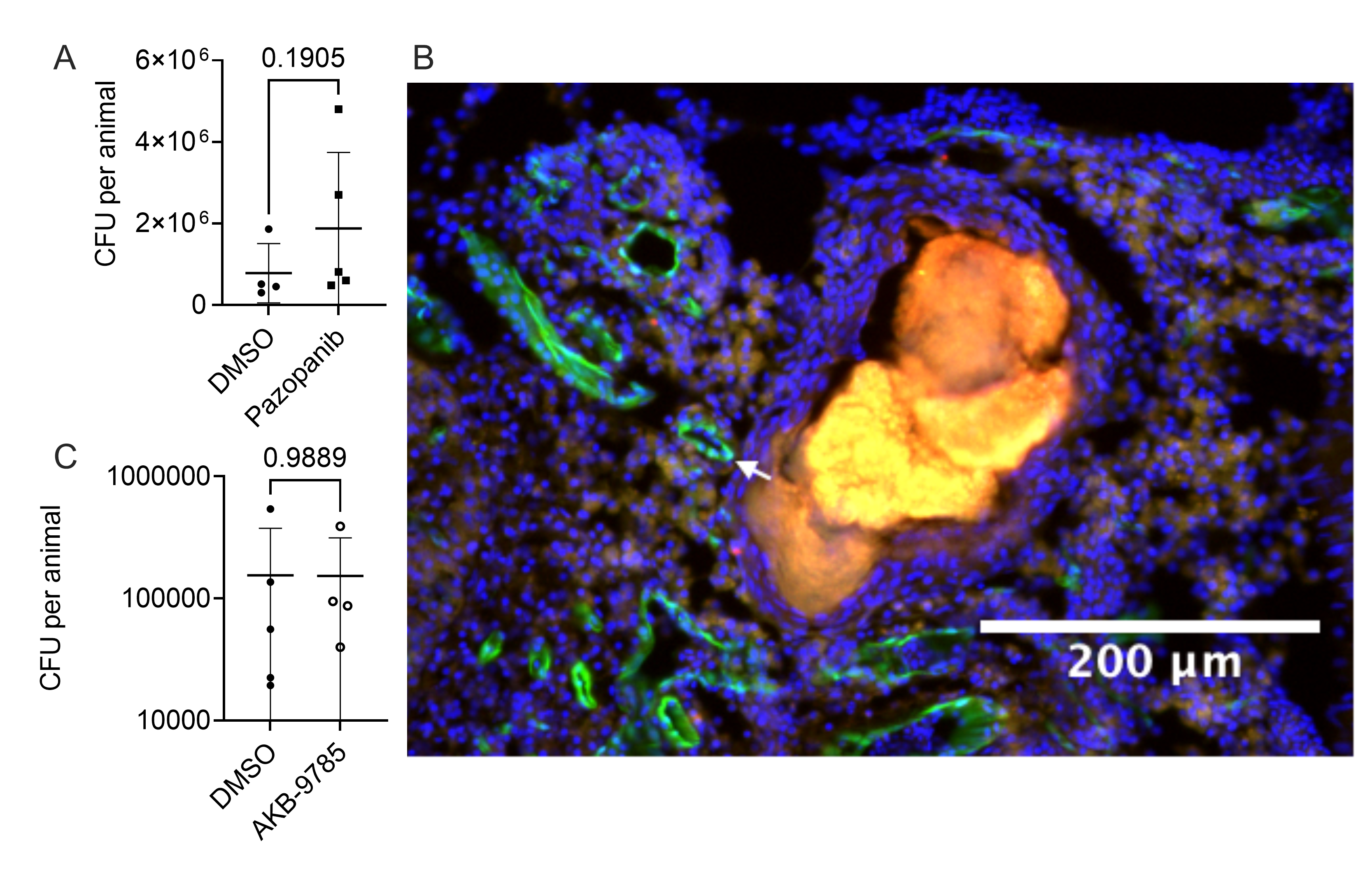
